# A framework for summarizing chromatin state annotations within and identifying differential annotations across groups of samples

**DOI:** 10.1101/2022.05.08.491094

**Authors:** Ha Vu, Zane Koch, Petko Fiziev, Jason Ernst

**Affiliations:** Bioinformatics Interdepartmental Program, University of California, Los Angeles, CA, 90095, USA; Department of Biological Chemistry, University of California, Los Angeles, Los Angeles, CA 90095, USA; Eli and Edythe Broad Center of Regenerative Medicine and Stem Cell Research at University of California, Los Angeles, Los Angeles, CA 90095, USA; Computer Science Department, University of California, Los Angeles, Los Angeles, CA 90095, USA; Jonsson Comprehensive Cancer Center, University of California, Los Angeles, Los Angeles, CA 90095, USA; Molecular Biology Institute, University of California, Los Angeles, Los Angeles, CA 90095, USA; Computational Medicine Department, University of California, Los Angeles, Los Angeles, CA 90095, USA; Illumina Inc

## Abstract

**Motivation:** Genome-wide maps of epigenetic modifications are powerful resources for non-coding genome annotation. Maps of multiple epigenetics marks have been integrated into cell or tissue type-specific chromatin state annotations for many cell or tissue types. With the increasing availability of multiple chromatin state maps for biologically similar samples, there is a need for methods that can effectively summarize the information about chromatin state annotations within groups of samples and identify differences across groups of samples at a high resolution.

**Results:** We developed CSREP, which takes as input chromatin state annotations for a group of samples and then probabilistically estimates the state at each genomic position and derives a representative chromatin state map for the group. CSREP uses an ensemble of multi-class logistic regression classifiers to predict the chromatin state assignment of each sample given the state maps from all other samples. The difference of CSREP’s probability assignments for two groups can be used to identify genomic locations with differential chromatin state patterns.

Using groups of chromatin state maps of a diverse set of cell and tissue types, we demonstrate the advantages of using CSREP to summarize chromatin state maps and identify biologically relevant differences between groups at a high resolution.

**Availability and implementation:** The CSREP source code is openly available under http://github.com/ernstlab/csrep.

## Introduction

Genome-wide maps of chromatin marks such as histone modifications and variants provide valuable information for annotating non-coding genome features (Barski *et al*., 2007; Ernst *et al*., 2011; Zhu *et al*., 2013; Xie *et al*., 2013). Efforts by large consortia and individual labs have produced chromatin state maps for many cell and tissue types (Roadmap Epigenomics Consortium et al., 2015; Consortium, 2012; Zhu *et al*., 2013; Barski *et al*., 2007). A popular representation of such data is chromatin states defined by the combinatorial and spatial patterns of multiple marks, which are generated by methods such as ChromHMM and Segway (Libbrecht *et al*., 2021)(Ernst and Kellis, 2010, 2012; Hoffman *et al*., 2012), and correspond to diverse classes of genomic elements including various types of enhancers and promoters.

Chromatin state maps have been produced for hundreds of different biological samples. In many cases there are multiple samples representing similar cell and tissue types (Boix *et al*., 2021; Roadmap Epigenomics Consortium et al., 2015). In such cases, to simplify analyses and visualizations, it may be desirable to have a single chromatin state annotation that summarizes the annotations for all samples from each group. A straightforward approach to this task is to take the most frequent chromatin state assigned at each position across samples. However, when the number of samples per group is small or the number of states is large, such an approach can be particularly vulnerable to noise. Furthermore, such an approach does not consider additional information available about the different chromatin states. For example, if a location was assigned to three different states in three samples, the summary annotation among these three states based on the frequency-based method would be arbitrary. However, by leveraging information about the co-occurrence of state assignments genome-wide, there is additional information to predict the most likely chromatin state annotation for a new sample from the group.

A related challenge is to identify differences in chromatin state annotations between two groups at a high resolution and on a per-state basis. Methods such as ChromDiff, chromswitch, and EpiAlign (Yen and Kellis, 2015; Jessa and Kleinman, 2018; Ge *et al*., 2019) can identify chromatin state differences between samples, but only calculate a measure of difference for a broad domain (e.g. a gene body), encompassing a large number of genomic bins for which the states are defined. Additionally, EpiAlign and Chromswitch are designed to measure the difference in annotations for one user-input query region in each run, and are not designed to generate genome-wide output, which is our focus. Another approach, EpiCompare (He and Wang, 2017) presented an approach for identifying differential enhancer chromatin states across cell or tissue tissue types, but did not consider other types of chromatin states. SCIDDO (Ebert and Schulz, 2020) can detect genome-wide significant differential chromatin domains between two groups of samples while incorporating a measure of similarity among states. However, SCIDDO only provides a single differential score per position and does not directly answer the question of what chromatin state switch occurs at each genomic position. Another method, dPCA (Ji *et al*., 2013), works directly on chromatin mark signals and does not quantify state differences across groups of samples.

To effectively summarize the chromatin state annotations for a group of samples and prioritize the chromatin state differences between two groups on a per-state basis, at high resolution, we introduce CSREP. CSREP leverages both the information about the input samples’ chromatin states at a position being summarized, as well as information of states’ co-occurrences in different samples within the same group across the genome. CSREP does this by first generating probabilistic estimates of chromatin state annotations by using an ensemble of multi-class logistic regression classifiers that predict the state assignment in a sample at a position, given the annotations in other samples, at the corresponding genomic position. From those predictions, CSREP is then able to produce a single summary state assignment per position. CSREP can also use the difference of summary probabilistic predictions for two groups of samples to quantify the difference in state assignments between the two groups on a per-state basis, e.g. one genome-wide score track per chromatin state.

Using CSREP, we generated the summary chromatin state maps for 11 groups of tissue/cell types from Roadmap Epigenomic Project (Roadmap Epigenomics Consortium et al., 2015), and for 75 groups from the Epimap Portal (Boix *et al*., 2021). We show that CSREP can better predict chromatin state assignments in held-out samples than a counting-based baseline method. We also verify that the resulting summary chromatin maps show correspondence with the group’s average gene expression profile. Additionally, we show that CSREP’s differential scores can recover differential epigenetic signals on chromosome X between male and female samples. We also show that CSREP differential scores between samples from two different tissue groups can predict regions of differential peaks for various chromatin marks. The CSREP implementation is designed to be user-friendly and includes a detailed tutorial, available at https://github.com/ernstlab/csrep. We expect CSREP will be a useful tool for summarizing chromatin state maps within groups and finding differences across groups. Additionally, the summary annotations for different tissue groups that we generated with CSREP are expected to be useful resource.

## Results

### CSREP method

CSREP takes as input chromatin state maps for a group of samples learned based on a concatenation approach (Ernst and Kellis, 2010, 2012) to ensure that annotations for different samples share chromatin state definitions. CSREP then generates as output (1) a summary probabilistic chromatin state assignment matrix and (2) a summary state map track for the group. The summary state assignment matrix represents the probabilities of each state being present at each genomic position in a new sample of that group. To generate these, CSREP takes a supervised learning approach, leveraging information about the co-occurrence of states from the different samples across the genome. Specifically, for each group of input samples, CSREP trains an ensemble of multi-class logistic regression classifiers (Hastie *et al*., 2009) to generate probabilistic predictions for each chromatin state at each position **(Fig. 1A, Methods)**. We used multi-class logistic regression classifiers since they provide well calibrated probabilities, are robust, and relatively fast to train. Each classifier is trained with *labels* based on the chromatin state assignments from one sample and *features* based on the chromatin states in other samples for the same genomic positions. Each classifier then makes a probabilistic prediction of the chromatin state assigned at each genomic position in the target sample. The chromatin state input features to each logistic regression classifier are represented with a one-hot-encoding of the chromatin states. The classifiers are trained on randomly selected genomic positions that constitute 10% of the genome, while the predictions are calculated genome-wide. The resolution of predictions is the same as that of input samples’ chromatin state maps (200bp with default settings from ChromHMM). The prediction result for each sample’s chromatin state map are represented in a matrix with *rows* corresponding to genomic positions and *columns* chromatin states. The values in each row sum to 1, representing the probabilities of state assignments at a genomic position. The probabilistic summary of a group is based on averaging the prediction output matrices for each sample in the group. These probabilistic predictions are then used to generate a summary chromatin state map for the group of samples by assigning the state with maximum assignment probability to each genomic position.

**Fig. 1:**
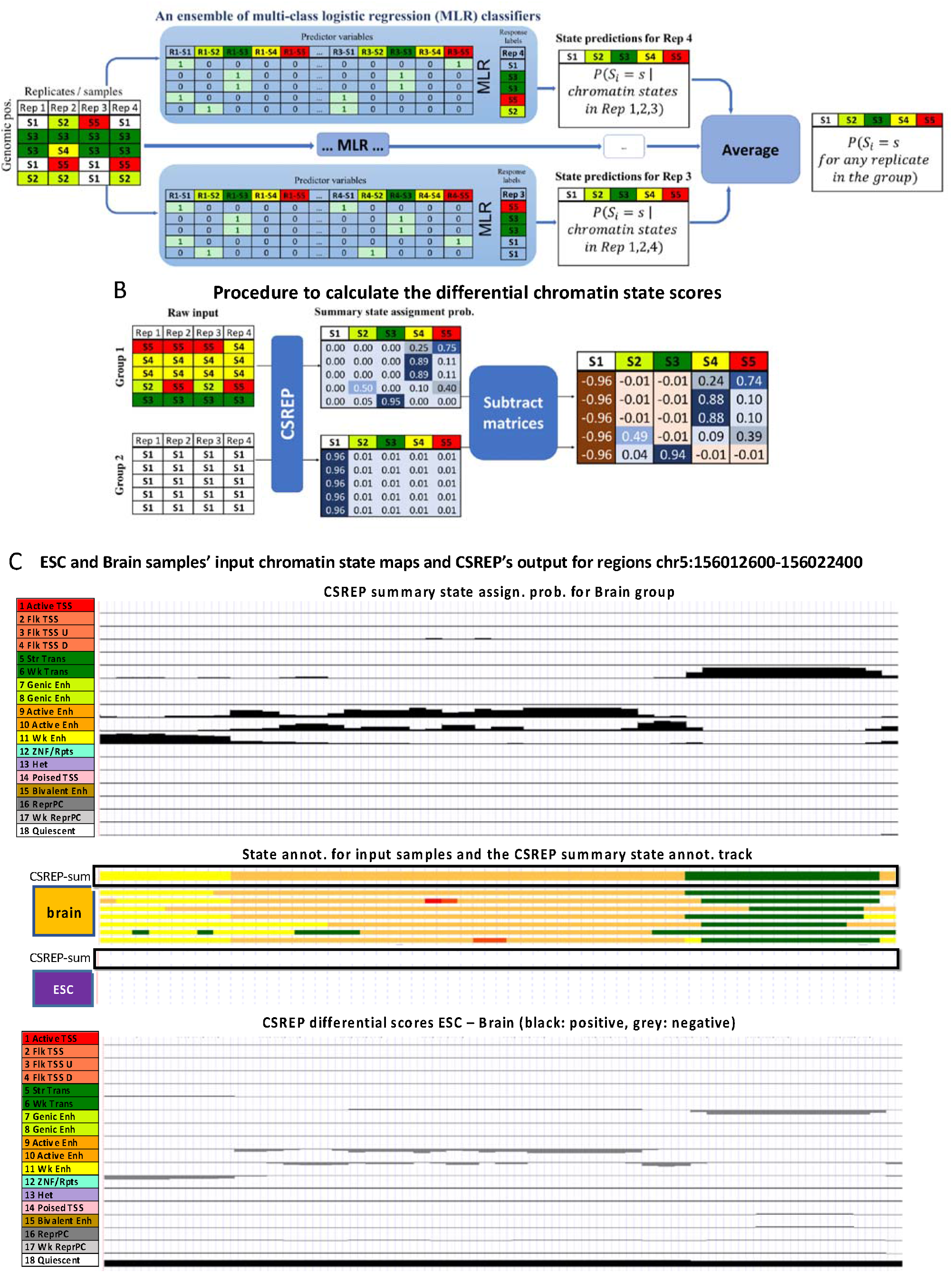
Overview of CSREP. **(A)** CSREP uses an ensemble of multi-class logistic regression models. In each model, the chromatin state map at the target sample is predicted based on the one-hot encoding of chromatin state assignments at the corresponding genomic positions in other samples. Multi-class logistic regression outputs the probabilities that each genomic position (row) in the target sample will be assigned to each state (column). CSREP averages the prediction matrices for target samples, to output the summary state assignment probability matrix. **(B)** The operations to obtain differential chromatin state assignment scores between two groups with multiple samples. CSREP calculates the summary chromatin state assignment matrices for two groups, and subtracts one group’s summary matrix from the other’s to obtain differential chromatin scores. Different chromatin scores are bounded between -1 (brown) and 1 (blue). **(C)** Visualization of CSREP’s output in a genomic region (hg19, chr5:156,012,600-156,022,400). The top of the subpanel shows the CSREP’s summary chromatin state probabilities for 18 states across seven Brain reference epigenomes. Each track shows the probabilities of assignment for one state, as named and colored on the left. The middle subpanel shows the 18-state chromatin state maps for 7 Brain samples and 5 ESC samples from Roadmap Epigenomics (Roadmap Epigenomics Consortium et al., 2015), and the CSREP’s output summary chromatin state maps for each group, outlined in black. States are colored as in legend as at the top of this subpanel. The last subpanel shows the differential chromatin scores when ESC’s summary state probabilities are subtracted from Brain’s. Each track shows one state’s differential scores. Scores between 0 and 1 are colored black, while those between -1 and 0 are colored grey.

CSREP’s summary probabilistic predictions can be directly used to generate differential chromatin state maps for two groups with multiple samples. This is achieved by subtracting the summary chromatin state assignment matrices of one group (first group) from the other’s (second group) **(Fig. 1B, Methods)**. At each genomic position, CSREP’s chromatin differential scores for individual chromatin states are bounded between -1 and 1, with a score of 1 in state *S* meaning state *S* was predicted to be the annotation for the first and second groups with probabilities 1 and 0 respectively, and vice versa for -1 (**Fig. 1C**). Overall, in addition to summarizing the state assignments for groups of samples, CSREP can calculate scores of differential chromatin state assignments for pairs of groups at the resolution of the input chromatin state maps.

### CSREP is predictive of chromatin states on held-out samples

We applied CSREP to a compendium of 18-state chromatin state maps for 64 samples (reference epigenomes) from 11 tissue groups generated by the Roadmap Epigenomics Project (Roadmap Epigenomics Consortium et al., 2015). The tissue groups include embryonic stem cells (ESCs), induced pluripotent stem cells (iPSC), ESC-derived cells, blood & T-cells, HSC & B-cells, epithelial, brain, muscle, heart, smooth muscle and digestive. The numbers of input samples for each tissue group range from 3 to 12. We provide the CSREP’s genomewide summary probabilistic and hard state assignments for 11 tissue groups (**Data availability**).

We first visualized CSREP’s summary chromatin state maps for a group of samples from digestive and heart tissue groups, which have 10 and 3 samples, respectively **(Fig. 2A, Supp. Fig. 1-4)**. For each group, we arbitrarily selected four 500-kb regions and visualized the input chromatin state maps and CSREP’s output probabilistic estimates and summary state map at such genomic windows. We observed expected correspondence between the groups’ input and output chromatin state assignment estimates **(Fig. 2A, Supp. Fig. 1-4)**.

**Fig. 2:**
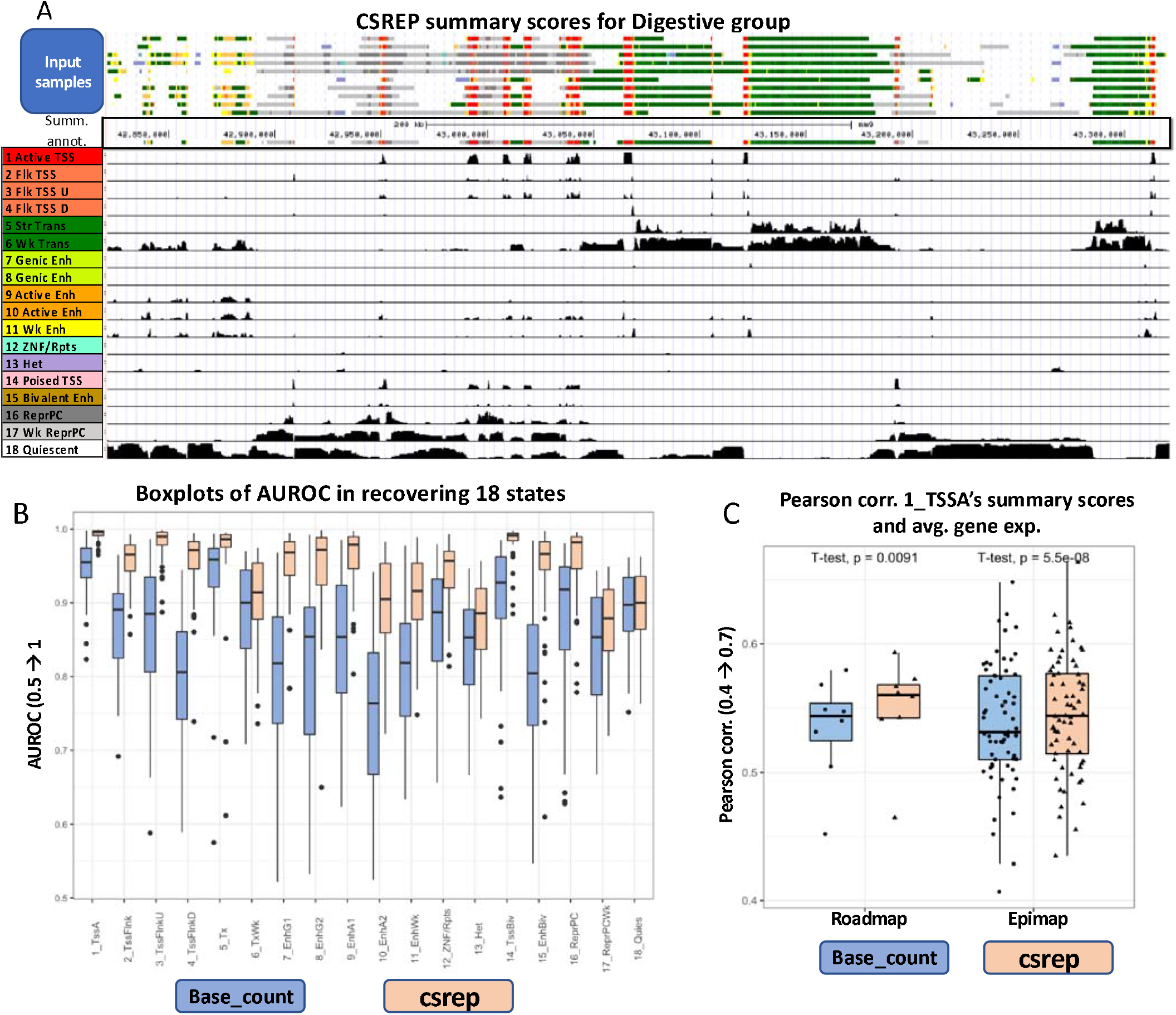
Performance of CSREP in summarizing a group with multiple samples’ chromatin state maps. **(A)** Visualization of one arbitrarily selected 500-kb region (chr5: 42,821,109-43,321,109, hg19). The first 10 tracks show chromatin state maps of 10 input samples from the Roadmap Epigenomics Consortium of the Digestive group, which were input to CSREP. The following track shows the summary chromatin state map from CSREP, which shows strong agreement with the input. States are colored based on the legend on the lower left. In the following 18 tracks, each track shows CSREP’s probabilities of assignment for each of 18 states, with state annotations in legend on left. **(B)** Boxplots showing the CSREP and base_count methods’ average, range and 25, 75% quantiles of the AUROCs across 64 samples, for each of the 18 chromatin states. The AUROCs were calculated in leave-one-out cross validation analysis where we used a group’s summary probabilistic chromatin state map to predict genomic locations of individual chromatin states in a left-out sample from the same cell/tissue group (**Supp. Methods**). States 1-18 (x-axis) are annotated as in **(A)**. **(C)** Boxplots showing the Pearson correlations between a group of samples’ (1) summary probabilities of state 1_TssA (active TSS) at annotated TSSs, and (2) the corresponding groups’ average gene expression (**Supp. Methods**). We obtained the correlations for 11 groups of cell types from the Roadmap Epigenomics Project, and 65 groups from EpiMap. Each dot shows the Pearson correlation for data from a group of samples.

To quantitatively evaluate CSREP’s summary output for a group of samples, we evaluated the accuracy of CSREP’s summary probabilistic chromatin state predictions in a leave-one-out cross-validation analysis. In particular, for each chromatin state, we calculated Area Under the Receiver Operating Characteristic (AUROC) curve for predicting genomic locations assigned to the state in the left-out sample from the group **(Supp. Methods)**. We compared the performance of CSREP against a baseline method, denoted base_count, which counts each state’s frequency across input samples at each genomic position (**Supp. Methods**).

CSREP showed strong predictive performance for chromatin states in left-out samples with average AUROCs across 64 samples varying from 0.871 to 0.993 for the 18 states. Across the 18 states, CSREP consistently had better AUROC in recovering individual states compared to the baseline method base_count **(Fig. 2B)**. The average AUROC improvements by CSREP compared to base_count ranged from 0.003 (for state 18_Quies) to 0.157 (for state 4_ TSSFlnkD). Larger performance improvements by CSREP relative to base_count are observed for all chromatin states when there are fewer input samples in the group (**Supp. Fig. 5**).

### CSREP summary chromatin state maps’ association with gene expression

Transcription start sites (TSS) are marked by different histone modifications and variants that can correlate gene transcription (Kimura, 2013; Soboleva *et al*., 2014). Here, we evaluated how CSREP’s summary state map for a tissue group is predictive of the group’s gene expression profiles at transcription start sites (TSS) of genes. First, we obtained gene expression data for available samples for the 11 tissue groups as above, and calculated the average protein-coding gene expression for each group (**Supp. Methods**). We then calculated the Pearson correlation between (1) the group’s average expression for protein coding genes and (2) CSREP’s summary state assignment probabilities for state 1_TssA (active TSS state) at the corresponding genes’ TSSs. We did the same evaluation for base_count. CSREP had significantly higher correlations than base_count (paired t-test p-value: 0.009, average 0.550 vs. 0.534, **Supp. Methods**). We next extended this analysis for a larger dataset for 552 samples in 75 groups from EpiMap repository based on state 1_TssA from the same 18-state annotations (Boix *et al*., 2021) (**Supp. Methods**). The 75 groups were previously formed based on tissue types and developmental stages with the number of samples per group ranging from 3 to 38 (**Supp. Methods, Data Availability**). Of the 75 groups, 65 also had gene expression data available for at least one sample. Across these 65 groups, again CSREP’s had significantly higher correlations than base_count (paired t-test p-val: 5.5e-08, average 0.545 vs. 0.538, **Supp. Methods**). Overall, CSREP’s summary chromatin state maps at TSS for the TssA state show significantly higher correspondence with gene expression levels compared to the base_count method.

### CSREP detects differential chromatin regions associated with different sexes

We next investigated the performance of CSREP at identifying biologically meaningful chromatin state changes between groups of male and female samples based on its ability to prioritize chromatin state differences on chromosome X (chrX) relative to autosomal chromosomes. Specifically, we applied CSREP to calculate differential chromatin state scores between 25 female and 44 male samples from Roadmap Epigenomics (**Supp. Methods**) (Yen and Kellis, 2015; Ge *et al*., 2019) by subtracting CSREP’s summary state probability matrix for the female samples from the corresponding matrix for the male samples.

We analyzed CSREP’s differential scores for all chromatin states across autosomal chromosomes and chrX (**Fig. 3A, Supp. Fig 6-7**). Three states with the largest magnitude of difference in mean scores between the sex chrX and autosomes were states 13_Het (heterochromatin, marked by H3K9me3), 17_ReprPCWk (weak polycomb repressed complex) and 18_Quies (quiescent). In chrX, compared to autosomal chromosomes, the distribution of differential scores for states 13_Het and 17_ReprPCWk showed a larger tail of negative. ChrX’s average score minus the autosomes’ average score values for states 13_Het and 17_ReprPCWk were -0.039 and -0.054, respectively (**Supp. Fig. 7**), implying that on chromosome X, female samples are more often assigned to these states compared to male samples. State 18_Quies showed the opposite trend with a difference of 0.11(**Fig. 3A, Supp. Fig. 7**). These results are consistent with sex-specific chrX inactivation, which is used in female mammals to achieve dosage compensation between the two sexes (Wutz, 2011; Yen and Kellis, 2015).

**Fig. 3:**
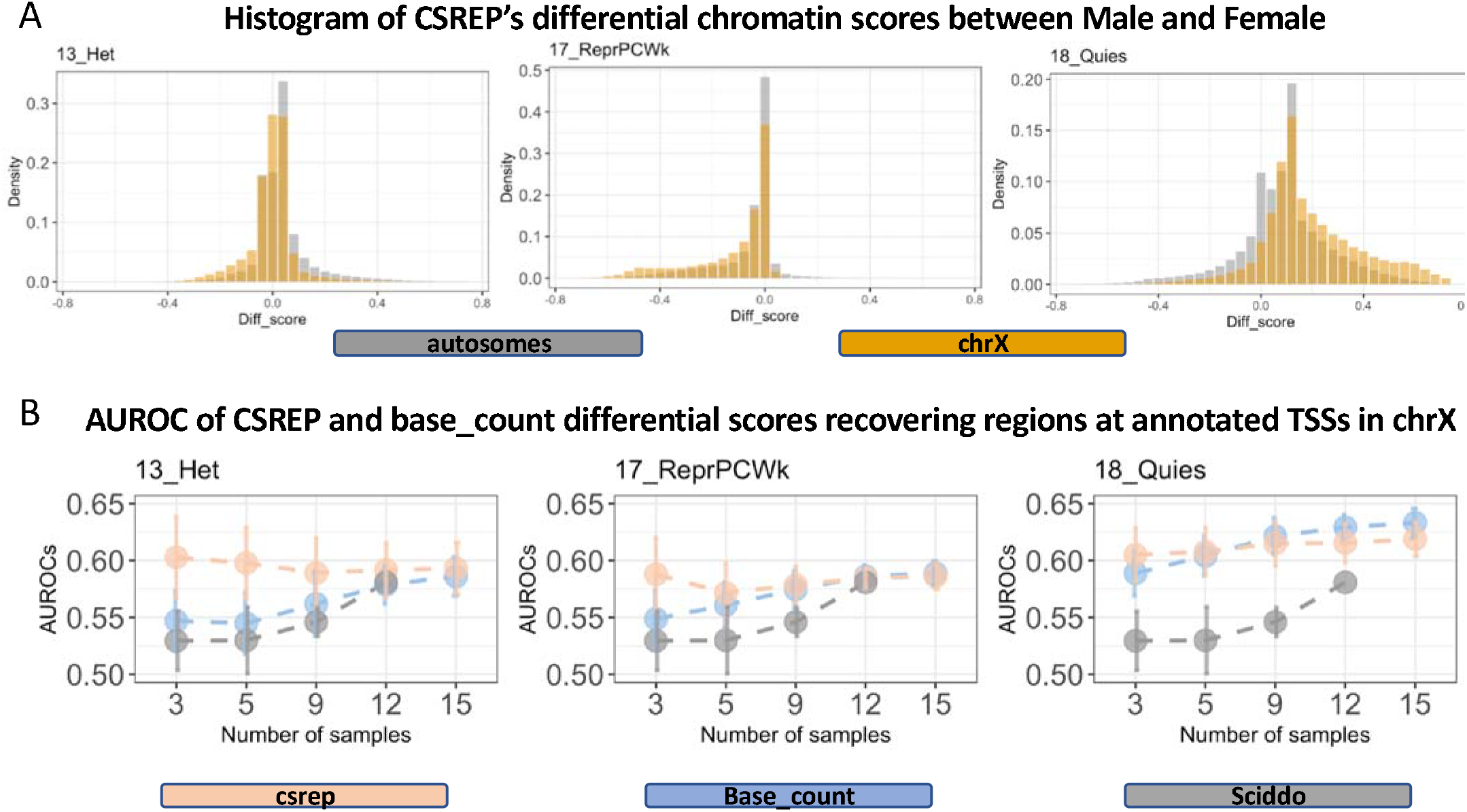
CSREP show signals of differential chromatin state scores in chromosome X when comparing male and female samples. (A) Each subpanel shows the histogram of CSREP’s differential scores in autosomes and chromosome X, for states associated with heterochromatin (13_Het), weak polycomb repressed domains (17_ReprPCWk), and quiescent regions (18_Quies). The x-axis shows differential scores, with positive values implying male samples have higher probabilities of being in the state compared to female samples, and vice versa for negative values. Histograms of scores for all states are in **Supp. Fig 6. (B)** AUROCs of recovering regions overlapping annotated TSSs on chromosome X, using differential chromatin scores of three states as in **(A)**, outputted by CSREP and base_count for Male and Female groups (**Supp. Methods)**. We calculated the AUROCs using different sets of input male and female samples, with varying number of samples in each group (x-axis). For each number of samples (x-axis), we conducted the analysis for 30 sets of male and female input samples (**Supp. Methods**). The plots show the average (dots) and standard deviation (error bars) of the AUROCs across the 30 sets of input samples. SCIDDO did not successfully generate output for the case of 15 input samples so no results are reported for that.

We next compared the performance of CSREP and other methods in recovering annotated transcription start sites (TSSs) on chrX, using the above-mentioned states, given varying numbers of input samples (**Supp. methods**) (**Fig. 3B**). To do this, we randomly selected 30 subsets of size *n* male and *n* female samples from the set of available 44 male and 25 female samples, where *n* is varied within the set of 3, 5, 9, 12 or 15 samples. Given each set of input male and female samples, we calculated the receiver operating characteristic (ROC) curve when using differential chromatin scores between male and female groups to predict locations overlapping annotated TSSs on chrX, against the background of those overlapping all annotated TSSs in the genome (**Supp. Methods**). We observed that CSREP showed the largest advantage over base_count, as measured by AUROCs, when the number of input samples from Male and Female groups is relatively small, e.g. 3 samples in each group (**Fig. 3B**). As the number of input samples from each group increases sufficiently, the overall gap of performance between CSREP and base_count goes away. In all cases, CSREP and base_count show better performance compared to SCIDDO (Ebert and Schulz, 2020) (**Fig. 3B**). Overall, CSREP showed the greatest advantage over other approaches when the number of samples is relatively small, which occurs frequently in practice.

### CSREP differential scores recover differential chromatin mark peaks

We next analyzed how well CSREP’s, base-count’s and SCIDDO’s differential chromatin state scores can predict genomic regions overlapping differential signals of DNase I hypersensitivity (DNase), H3K9ac and H3K27ac between samples from embryonic stem cell (ESC) and brain. DNase and H3K9ac signals were not used for learning the 18-state model used to annotate the two groups’ input samples, providing an independent validation. While H3K27ac was used in learning the input chromatin state maps, since all the methods being compared (CSREP, base_count, SCIDDO) had access to the same maps as input, and H3K27ac is a well-established mark of cell-type specific activity (Creyghton *et al*., 2010), we still considered H3K27ac in the evaluations of methods’ performance.

For each of the three chromatin marks, we first obtained a set of bases that are present in peaks in all samples from ESC but not in any from the Brain group and vice versa (**Supp. Methods**). We then calculated CSREP and base_count differential chromatin scores by subtracting the summary chromatin state map of Brain from that of the ESC. Additionally, we applied SCIDDO to the same set of input data (**Supp. Methods**). We evaluated, in terms of AUROC, how well the methods prioritize regions overlapping bases in the ESC-/brain-specific sets of peaks (**Supp. Methods**). For CSREP and base_count, we conducted separate evaluations for each chromatin state, but did *not* for SCIDDO since it outputs one score track that measures the overall difference across the chromatin state landscape between the two groups.

Across the different marks being evaluated, the highest AUROCs were consistently from CSREP based on its scores for either from promoter or enhancer associated states (**Fig. 4**). For example, for identifying brain specific H3K9ac peaks, CSREP had an AUROC of 0.717 based on the evaluation with state 9_EnhA1, an active enhancer state, while the maximum AUROC achieved for base_count was 0.617 and SCIDDO’s AUROC was 0.564. These analyses suggest that CSREP differential scores better correspond to locations of individual mark differences between two groups of samples genomewide, compared to other approaches that also aim to identify chromatin state differences between two groups. The advantage of CSREP over SCIDDO may in part be due to CSREP producing scores with respect to specific chromatin states and including the direction of change (with positive/negative scores implying one group’s higher state assignment probabilities compared to the other’s).

**Fig. 4:**
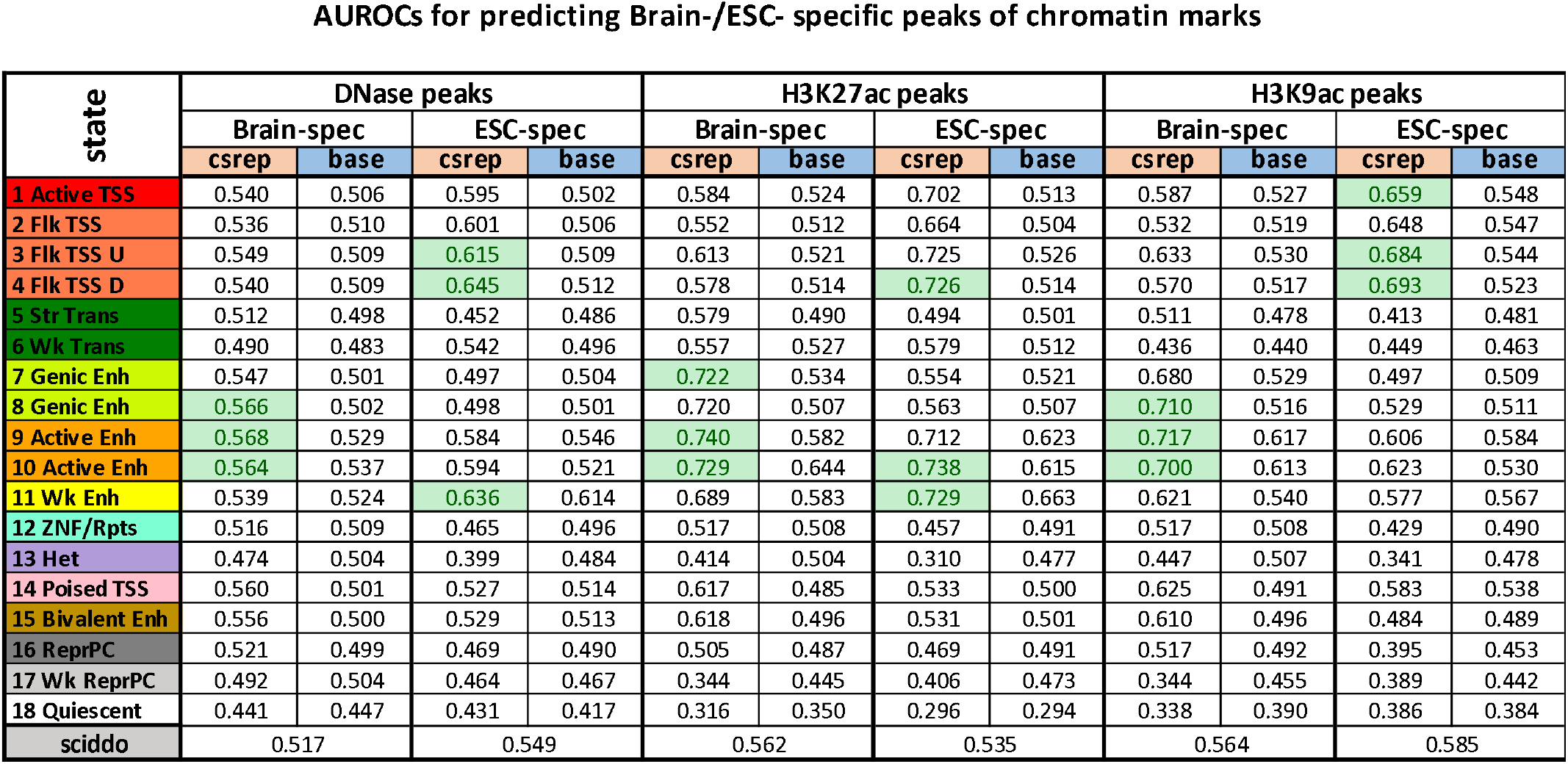
CSREP better recovers differential chromatin marks signals between ESC and Brain. The table shows AUROCs for differential scores’ predictions of genomic regions associated with differential peak signals for one chromatin mark, from left to right: DNase, H3K27ac and H3K9ac. For each chromatin mark, it shows the AUROCs of predicting signal peaks observed in Brain and ESC exclusively (Brain-spec and ESC-spec). Differential scores outputted by CSREP or baseline are shown for each chromatin state (rows). In each category of comparisons, the top three scores that show highest AUROCs are highlighted in green. Along the bottom is the AUROC for SCIDDO.

## Discussion

Here, we proposed CSREP, a method for probabilistically summarizing the chromatin state maps from a group of samples. CSREP achieves this by training multi-class logistic regression models to predict the chromatin state annotations of one sample using data from others, and then averaging the prediction probabilities across all samples in the group. CSREP outputs the probabilities of each chromatin state being assigned to each genomic position, at the same resolution that chromatin states are annotated. We applied CSREP to generate summary 18-state chromatin state assignment probability matrices for 11 groups of cell and tissue types from Roadmap Epigenomics Project (Roadmap Epigenomics Consortium et al., 2015), and 75 groups of samples stratified by cell and tissue types and developmental phases from EpiMap (Boix *et al*., 2021), and have made them publicly available (**Data Availability**).

Our analyses reveal that CSREP’s probabilistic summary of state assignments better predicts the chromatin states of held out samples compared to the counting-based baseline approach. We also showed that CSREP’s summary assignment probabilities of state 1_TssA at TSS was well correlated with the average gene expression of the group, and significantly higher than those achieved by the counting-based baseline.

CSREP can also be used to directly quantify the difference in chromatin state maps between two groups with multiple samples, at the resolution of the input annotations. CSREP produces differential scores for each chromatin state at each genomic position, which represent the difference in probabilities that samples from two input groups are assigned to each specific state. Therefore, CSREP differential scores are bounded (−1 to 1), interpretable with respect to specific chromatin state changes, and indicative of the direction of change, which contrasts it with other approaches that provide a single score showing magnitude of difference per genomic position. We used CSREP to compare the chromatin state annotations between male and female samples from Roadmap Epigenomics (Roadmap Epigenomics Consortium et al., 2015), and showed that CSREP can better predict regions overlapping genes’ TSS on chrX, particularly when there are few samples in each group. CSREP’s differential scores for states associated with active enhancers and promoters better recovered tissue-group-specific peaks of DNase/H3K27ac/H3K9ac signals compared to alternative approaches, suggesting that CSREP provides useful additional information for analyzing epigenomic changes across tissue types. Future work could apply CSREP to compare additional biological conditions or disease state (e.g. cancer vs non-cancer).

CSREP works directly off of chromatin state annotations, which makes CSREP agnostic to the specific methods used to produce those annotations. Some methods for learning chromatin state annotations have the option to expose posterior probability estimates of annotations, which could potentially be used in an extended version of CSREP. However, assuming accurately determined posterior probability estimates are available as input would also make CSREP less generally applicable.

To facilitate the use of CSREP, we provide an implementation of CSREP as a snakemake pipeline (Mölder *et al*., 2021) with a detailed tutorial that only requires users to modify parameters in a yaml file. The program can be run either on local computers or on computing clusters, in which case snakemake will optimize the workflow for execution.

We expect CSREP to be a useful tool and the output we have provided from it a valuable resource for summarizing summarize chromatin state maps from groups of samples and prioritizing regions with differential chromatin state changes across pairs of groups of samples.

## Methods

### CSREP’s summarization of a group of samples

Let *G* denote the number of genomic bins across the genome, *S* the number of chromatin states, and *N* the number of samples in the target group of samples. Let *C*_*i,n*_ denote the chromatin state assigned to sample *n* at genomic position *i*, which can take one value of 1, 2, …, *S*. Let *N*_*n*_ denote the set of samples not including *n*, i.e. *N*_*n*_ = {1, …, *N*} − {*n*}. In general, CSREP is an ensemble of *N* multi-class logistic regression classifiers such that for each sample *n*, CSREP trains a classifier to predict the chromatin state map of this sample based on features in the remaining samples (*N*_*n*_). The predictor variables for such a model include one-hot encoding chromatin state maps of the *N* − 1 samples (all samples in the group except *n*) and an intercept term, resulting in (*N* − 1) \**S* +1 predictor variables. The response variable is the chromatin state of the target sample n, which can take one value of 1, 2,.. ., *S*.

In the multi-class logistic regression model, let *X*_*i*_ denote the vector of predictor variables at position *i*, which has length (*N*-1)\**S* +1 and takes values {0,1}. The last entry of *X*_*i*_ is 1, corresponding to the intercept term. Let *Y*_*i*_ denote the value of the response variable at position *i*, which takes values {1,2, …, *S*}. Since the input chromatin state maps segment the genome into 200-bp bins, we refer to each genomic position as one 200-bp window in the genome. We randomly selected genomic positions for the training data set, such that these positions constitute 10% of the genome. Given the training data set, for each state *s* ∈ {1, …, *S* − 1}, the multi-class logistic regression model learns a coefficient vector *β*_*s*_ with length (*N* − 1)\**S* + 1, corresponding to the number of predictor variables. The probability of sample *n*’s chromatin state s being assigned at position *i* is calculated as:

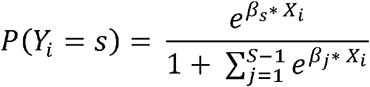

for *s* ∈ {1, . ., *S* − 1}, and as the following when *s* = *S*:

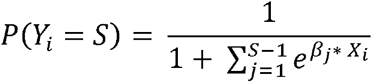

After CSREP trains the multi-class logistic regression model on training data that constitute 10% of the genome, and *l*2-norm penalty. The model is implemented using Python’s sklearn, pybedtools package and snakemake (Dale *et al*., 2011; Quinlan and Hall, 2010; Mölder *et al*., 2021). CSREP applies the model to generate predictions of genome-wide probabilistic chromatin state map for sample *n*, which is presented in a matrix of size *G* * *S*. The output matrices from *N* predictions for *N* samples are then averaged, so at each genomic bin, the sum of state assignment probabilities across *S* states is 1. In addition, the chromatin state with the maximum probability in each row is recorded to produce a single representative chromatin state map for the entire group of samples.

### CSREP’s application to prioritizing differential chromatin state changes between two groups of samples

To calculate differential chromatin state maps between two groups of samples, group1 and group2, CSREP first calculates the probabilistic chromatin state map matrices for each group as described above, denoted as *R*_1_ and *R*_2_, respectively. After this, CSREP subtracts the two matrices to represent the differential chromatin state map between group1 and group2 (denoted *D*_12_), i.e. *D*_12_ = *R*_1_ − *R*_2_. We note that we used signed and not absolute difference here and thus the score range from − 1 to 1. A score on row i and column *s* of *D*_12_, denoted *D*_12,*i,s*_, being − 1 means group2 is estimated to have probability 1 of being assigned to state *s* at position *i* while group1 has probability of 0. Additionally, since CSREP assigns *S* scores of differential chromatin maps to each genomic position *i*, corresponding to *S* states, CSREP can uncover specific chromatin states switch. For example, if *D*_12,*i,s*_ =0.8 when *s* = 1 while *D*_12,*i,s*_ = − 0.8 when *s*= 2, we can say it is likely that at position *i*, group1 is more likely to be in state 1 while group2 is likely to be in state 2.

## Supporting information

AF2_metadata_sample

## Data availability

The summary chromatin state maps (the chromatin state assignment matrices and the corresponding state annotation) for 11 tissue groups in Roadmap Project and 75 groups in Epimap Portal are available for download at https://github.com/ernstlab/csrep. The summary state maps for samples in Roadmap Epigenomics and EpiMap are provided both in hg38 and in hg19.

## Acknowledgements

We thank all members of the Ernst lab for their helpful suggestions on the manuscript.

## References

Barski, A. et al. (2007) High-resolution profiling of histone methylations in the human genome. Cell, 129, 823–837.

Boix, C.A. et al. (2021) Regulatory genomic circuitry of human disease loci by integrative epigenomics. Nature, 590, 300–307.

Consortium, E.P. (2012) An integrated encyclopedia of DNA elements in the human genome. Nature, 489, 57–74.

Creyghton, M.P. et al. (2010) Histone H3K27ac separates active from poised enhancers and predicts developmental state. Proc. Natl. Acad. Sci., 107, 21931–21936.

Dale, R.K. et al. (2011) Pybedtools: a flexible Python library for manipulating genomic datasets and annotations. Bioinformatics, 27, 3423–3424.

Ebert, P. and Schulz, M.H. (2020) Fast detection of differential chromatin domains with SCIDDO. Bioinformatics.

Ernst, J. et al. (2011) Mapping and analysis of chromatin state dynamics in nine human cell types. Nature, 473, 43–49.

Ernst, J. and Kellis, M. (2012) ChromHMM: automating chromatin-state discovery and characterization. Nat. Methods, 9, 215–216.

Ernst, J. and Kellis, M. (2010) Discovery and characterization of chromatin states for systematic annotation of the human genome. Nat. Biotechnol., 28, 817–825.

Ge, X. et al. (2019) EpiAlign: an alignment-based bioinformatic tool for comparing chromatin state sequences. Nucleic Acids Res., 47, e77–e77.

Hastie, T. et al. (2009) The elements of statistical learning: data mining, inference, and prediction Springer.

He, Y. and Wang, T. (2017) EpiCompare: an online tool to define and explore genomic regions with tissue or cell type-specific epigenomic features. Bioinformatics, 33, 3268–3275.

Hoffman, M.M. et al. (2012) Unsupervised pattern discovery in human chromatin structure through genomic segmentation. Nat. Methods, 9, 473.

Jessa, S. and Kleinman, C.L. (2018) Chromswitch: a flexible method to detect chromatin state switches. Bioinformatics, 34, 2286–2288.

Ji, H. et al. (2013) Differential principal component analysis of ChIP-seq. Proc. Natl. Acad. Sci., 110, 6789–6794.

Kimura, H. (2013) Histone modifications for human epigenome analysis. J. Hum. Genet., 58, 439–445.

Roadmap Epigenomics Consortium et al. (2015)Integrative analysis of 111 reference human epigenomes. Nature, 518, 317–330.

Libbrecht, M.W. et al. (2021) Segmentation and genome annotation algorithms for identifying chromatin state and other genomic patterns. PLoS Comput. Biol., 17, e1009423.

Mölder, F. et al. (2021) Sustainable data analysis with Snakemake. F1000Research, 10.

Quinlan, A.R. and Hall, I.M. (2010) BEDTools: a flexible suite of utilities for comparing genomic features. Bioinformatics, 26, 841–842.

Soboleva, T.A. et al. (2014) Histone variants at the transcription start-site. Trends Genet., 30, 199–209.

Wutz, A. (2011) Gene silencing in X-chromosome inactivation: advances in understanding facultative heterochromatin formation. Nat. Rev. Genet., 12, 542–553.

Xie, W. et al. (2013) Epigenomic analysis of multilineage differentiation of human embryonic stem cells. Cell, 153, 1134–1148.

Yen, A. and Kellis, M. (2015) Systematic chromatin state comparison of epigenomes associated with diverse properties including sex and tissue type. Nat. Commun., 6, 1–13.

Zhu, J. et al. (2013) Genome-wide chromatin state transitions associated with developmental and environmental cues. Cell, 152, 642–654.

