## Supplementary material for "A framework for summarizing chromatin state annotations within and identifying differential annotations across groups of samples": AF1_supp_materials

**Primary data sources**

We analyzed genome-wide 18-state chromatin state annotation for 64 reference epigenomes from the Roadmap Epigenomic Project Portal (Roadmap Epigenomics Consortium *et al.*, 2015) and 552 from the EpiMap Portal (Boix *et al.*, 2021). We will refer to each reference epigenome as a sample. State annotations data for samples from Roadmap were in hg19, while those from EpiMap were downloaded both in hg19 and hg38. The 18-state model was shared between Roadmap Epigenomics and EpiMap, and was trained based on data of 6 chromatin marks: H3K4me1, H3K4me3, H3K27ac, H3K27me3, H3K36me3 and H3K9me3. We assigned 64 samples from Roadmap Epigenomics into 11 groups based on the accompanying metadata’s tissue group labels. These groups include Blood & T-cell, Brain, Digestive, embryonic stem cells (ESC), ES-deriv, Heart, induced pluripotent stem cells (iPSC), Muscle, Skin, smooth muscle (Sm_Muscle) and HSC & B-cell. We assigned the biosamples from EpiMap into 75 distinct groups based on the metadata corresponding to unique combination of: extended biosample summary (tissue and sub-tissue types) and life stage (adult or embryonic, any biosamples with unknown life stage were filtered out from the analyses). We only analyzed groups of samples from EpiMap with at least 3 biosamples. Among sample groups from Roadmap Epigenomics, the number of samples per group ranged from 3 to 12, while the corresponding range for samples from EpiMap is 3 to 38. Details about the samples’ ID, groups and other metadata are provided in Additional File 2.

Using CSREP, we generated summary chromatin state maps for chromosomes 1-22 and X for the 11 groups from Roadmap Epigenomics using input data in hg19, and for the 75 groups from EpiMap using input data in both hg19 and hg38. We then used liftOver to lift the summary state maps for 11 groups from Roadmap Epigenomics from hg19 and hg38. This procedure first finds a one-to-one mapping for a subset of 200-bins between hg19 and hg38, i.e. if there are multiple bins from hg19 that got mapped to the same bin in hg38, those bins would not be included into the annotations. Then, we map both the state assignment probabilities and the summary state annotations for each bin in hg19 to the corresponding bin in hg38. Source code for the liftOver procedure, along with a detailed tutorial, is provided at <https://github.com/ernstlab/csrep>.

**Evaluation of CSREP in representing a group’s chromatin state maps**

We evaluated CSREP and an alternative baseline approach called base_count (defined below) for predicting representative chromatin state maps. We conducted this evaluation through a leave-one-out cross validation framework. Given a group with $N$ samples, for each sample indexed $n$, we evaluated the prediction of chromatin state map for sample $n$ when the state maps of the other $N-1$ samples are used as the input for generating the predictions. For these evaluations we used the data for the 64 samples from the 11 tissue groups from the Roadmap Epigenomics Project (Roadmap Epigenomics Consortium *et al.*, 2015) described above.

***Base_count method:*** Let $C_{nis} = 1$ if in sample $n$, at genomic position $i$, the observed chromatin state is $s$, and $C_{nis} = 0$ otherwise. The base_count approach represents the group’s chromatin state map by calculating the frequency of state $s$ being assigned at the $i$position (${BC}_{is}$) across the samples. In particular:

$${BC}_{is} = \frac{\sum_{n = 1}^{N} C_{nis}}{N}$$

Where *N* is the number of samples. Similar to CSREP, the output matrix for base_count method is of size $G * S$, with the sum of values in each row being 1. $G$ and $S$ represent the number of genomic bins the states, as explained in the main Methods.

***Calculating the ROC curves of prediction for a single chromatin state’s location:*** In each round of cross-validation, one sample with index $n$ is held-out. We then used CSREP and base_count to get the summary probabilistic chromatin state map for the group using input data from the remaining $N-1$ samples. For each state $s$, CSREP and base_count output the summary probability that each 200-bp genomic bin gets assigned to the state $s$. We divided the $[0,1]$probability range into 500 equal-width windows with lower bounds $l\in\{0, 0.002, ..., 0.998\}$. Within each probability window, any genomic positions with assignment probability for the state $s$ being no less than the window lower bound ($l$) will be predicted as being in state $s$ for sample $n$. Given the true chromatin state map in sample $n$, we then calculated the cumulative true positive rates and false positive rates of the prediction at each probability threshold $l$ to obtain the ROC curve. This analysis is repeated for each chromatin state, resulting in $S$ ROC curves.

***Evaluating CSREP’s summary chromatin state maps’ association with gene expression***

We obtained data of gene expression from the Roadmap Epigenomics Consortium (Roadmap Epigenomics Consortium *et al.*, 2015), which was available as a matrix of values in RPKM (reads per kilobase million) for genes in a subset of the samples from https://egg2.wustl.edu/roadmap/data/byDataType/rna/expression/57epigenomes.RPKM.pc.gz. We obtained the accompanying gene annotation information from https://egg2.wustl.edu/roadmap/data/byDataType/rna/expression/Ensembl_v65.Gencode_v10.ENSG.gene_info.gz. We filtered out genes that are not annotated as protein-coding, and transformed the gene expression matrix by adding a pseudo-count of 1 to the RPKM counts, and then log-transforming the resulting values. We also only included genes that are on chromosomes 1-22 and X in hg19 for this analysis, resulting in 20,787 distinct TSSs whose associated gene expression was available. For each of the 11 groups of samples in Roadmap Epigenomics, we obtained the group’s average gene expression profile by averaging over the gene expression values across all available samples in the group, i.e. samples that are both in the group and among the samples whose expression data was available. Among the 11 groups, 8 groups (all except for groups HSC & B-cell, iPSC and Sm_Muscle) had available gene expression data from the available samples. We then calculated the Pearson correlation of the summary chromatin state assignment probabilities for the 1_TssA state at positions that overlap with the 20,787 annotated TSSs and their corresponding average gene expression for each group.

For EpiMap, we obtained data of quantile-normalized protein coding genes’ expression from

https://personal.broadinstitute.org/cboix/epimap/rnaseq_data/merged_qn_log2fpkm.pc.mtx.gz, which is available as values in log2(FPKM), along with data of samples’ ID and genes’ EnsemblID. We utilized the same genes’ annotation information as provided by Roadmap Epigenomics, and only included protein-coding genes on chromosomes 1-22 and X in hg19. This resulted in 18,543 distinct TSSs whose associated gene expression was available from EpiMap. Among the 75 groups, 10 groups (SMTH.Digestive, HSC.MPP, EYE.EMB, EPTH.BREAST.EPITH, CA.UCEC, CA.RCC, CA.MYELOMA, BRN.EMB.BRN, BRN.CAUD.NUC, BONE.EMB) had no available gene expression data. We followed the same procedure mentioned above for the remaining 65 groups to obtain the Pearson correlation between a group’s average gene expression and summary state assignment probabilities for the 1_TssA state.

We used a paired t-test to compare the correlations resulting from CSREP against those from base_count, with the alternative hypothesis that CSREP’s correlations with gene expression are higher than base_count’s.

**Application of SCIDDO**

To run SCIDDO, we followed the tutorial provided by the author on Github https://github.com/ptrebert/sciddo/blob/master/testdata/tutorial.md (Ebert and Schulz, 2020), and generated a list of differential chromatin domains between two conditions. SCIDDO’s output contained overlapping differential chromatin domains with different values of SCIDDO scores. We then averaged the SCIDDO differential scores for overlapping regions of output differential chromatin domains, we assigned 0 to genomic regions not reported in SCIDDO’s output, implying no differential signals in chromatin states between the two groups.

***Evaluating CSREP’s differential chromatin state maps between Male and Female groups in recovering chromosome X- associated genomic regions***

For evaluating chromosome state differences between Male and Female groups with respect to chromosome X and autosomes, we obtained data for samples whose sex is annotated as either male or female (not ‘Unknown’ or ‘Mixed’), according to the provided metadata for the 98 samples with 18-state chromatin state maps from Roadmap Epigenomic Project (Roadmap Epigenomics Consortium *et al.*, 2015). In total, there are 44 Male samples and 25 Female samples. We then generated 30 sets of 3 male samples (randomly chosen from 44 Male samples) and 3 female samples (randomly chosen from 25 Female samples). We calculated the differential chromatin scores for the 30 sets of samples using CSREP, base_count and SCIDDO (also see section *Application of SCIDDO*). At the same time, we obtained a list of annotated TSSs for *protein-coding* genes from the accompanying metadata of genes’ coordinates and strand provided by the Roadmap Epigenomics project. For each set of 3 male and 3 female samples, we then obtained the *absolute values* of differential chromatin scores for regions that overlap these TSSs, and divided the score range window ($\left[ 0, 1 \right]$for CSREP and base_count, $[0, maximum value]$ for SCIDDO) into 100 equal-width bins. We then applied the same procedure as outlined in the above section to obtain true-/false- positive rates and AUROCs in predictions of TSS-overlapping regions on chromosome X, among all genomic regions overlapping annotated TSSs on the autosomes and chrX. We repeated the same analysis for a total of 30 sets of $n$ male and $n$ female samples, with $n \in\{3,5,9,12,15\}$. We note that not all 30 rounds of application of SCIDDO to calculate differential chromatin scores ran successfully, due to software failure. In particular, all applications of SCIDDO with 30 input sets of 15 male and 15 female samples (n=15) failed. For such cases, we report the average AUROCs for only successful runs of SCIDDO (**Fig. 3B**).

***Evaluating CSREP’s differential chromatin state map in recovering regions associated with differential chromatin mark signals***

To evaluate recovering differential chromatin mark signals, we first for DNase, H3K9ac and H3K27ac, downloaded the available broad peaks for samples from the ESC and Brain groups from Roadmap Epigenomics Project at https://egg2.wustl.edu/roadmap/data/byFileType/peaks/consolidated/broadPeak. A full list of links to data used is provided in **Additional File 2**. For each of the three chromatin marks and each cell group (ESC or Brain), we used $bedtools intersect$ (Quinlan and Hall, 2010) to obtain a set of peaks that are shared across all samples in the respective cell group. We then used $bedtools subtract$ function to derive peaks that are present in ESC samples and missing in Brain samples, and vice versa. We treated these ESC-specific, Brain-specific chromatin peaks as the ground-truth for this analysis. The number of base pairs for each group range from 2,735,377 bp (ESC-specific H3K9ac peaks) to 85,995,111 bp (Brain-specific H3K27ac peaks).

We generated differential chromatin state scores for CSREP for the ESC and Brain groups for Roadmap Epigenomics, by subtracting their probabilistic chromatin state predictions for Brain from those for ESC. The resulting differential chromatin matrices from the CSREP and base_count methods are of size $G*S$, and denoted $D_{CSREP}$ and $D_{BC}$, respectively, where  $D_{CSREP,i, s}$ and $D_{BC, i,s}$ denotes the CSREP and base_count differential score for state $s$ at genomic position $i$, respectively. We also obtained SCIDDO scores, which measure genome-wide differential chromatin patterns between the two groups, as described in section below. We denote the genome-wide SCIDDO score vector as$D_{SCIDDO}$, and $D_{SCIDDO, i}$ as the score at genomic position $i$.

To use $D_{CSREP}$ or $D_{BC}$ to calculate the ROC for genome-wide prediction of bases associated with ESC-specific or Brain-specific chromatin marks’ peaks, we divided the score range window $\left[ -1, 1 \right]$ into 200 equal-width bins with lower bounds $l \in\{-1, -0.99, ..., 0.99\}$. To calculate ROCs for predicting bases in ESC-specific peaks, for each state $s$ and each differential score lower-bound $l$, we set genomic positions where the differential scores for state $s$ being greater than or equal to $l$ denoted $\{i:D_{--, i, s}\geq l\}$. We compared such predictions with the ground-truth peaks described above to obtain the true- and false-positive rates of prediction for state $s$. To calculate ROCs for predicting bases associated with DNase/H3K9ac/H3K27ac Brain-specific peaks, first, we reversed the sign of  $D_{CSREP}$and $D_{BC}$, resulting in differential score matrices where positive values for state $s$ at  position $i$ implies that the respective position $(i)$ has a higher probability of being in state $s$ in Brain compared to ESC. Then, we applied a similar procedure as outlined above to calculate CSREP’s and base_count’s ROCs. To calculate the ROC curves based on $D_{SCIDDO}$, we divided the observed SCIDDO scores into 100 equal-width bins, ranging from the minimum to maximum values of the scores across the genome. We then applied the same procedure as for CSREP and base_count scores in one state, as mentioned above.

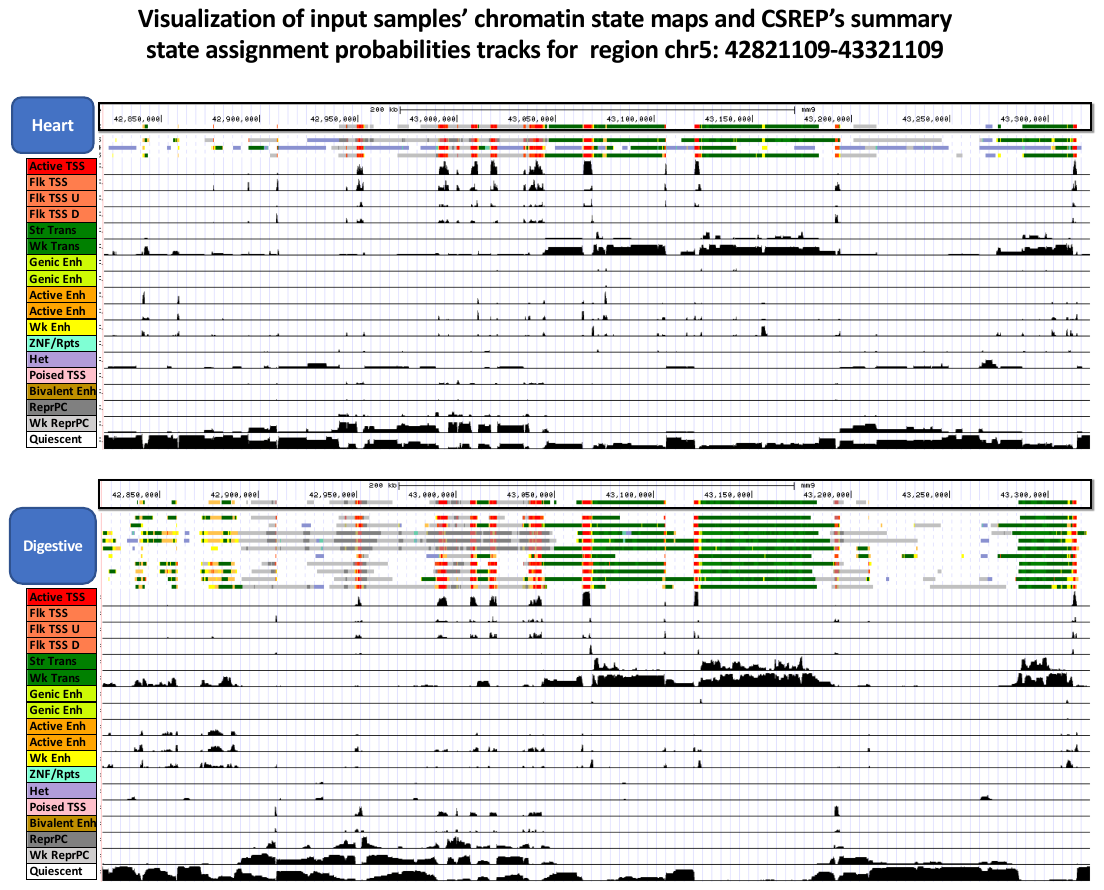

**Supplementary Figure 1: Visualization of CSREP’s input and output data for an arbitrary 500-kb genomic window (chr5: 42,821,109-43,321,109, hg19).** The visualization shows CSREP’s strong agreement with the chromatin state maps from 10 input samples from Digestive and 3 samples from Heart tissue groups from Roadmap Epigenomics Consortium. In each panel, the first track shows the summary chromatin state map based on CSREP. The following 3 (Heart) and 10 (Digestive) tracks show input samples’ chromatin state maps. States are colored based on legend on the left. In the following 18 tracks, each track shows the probabilities of assignment for one of 18 states. This region is the same as in **Fig. 2A**.

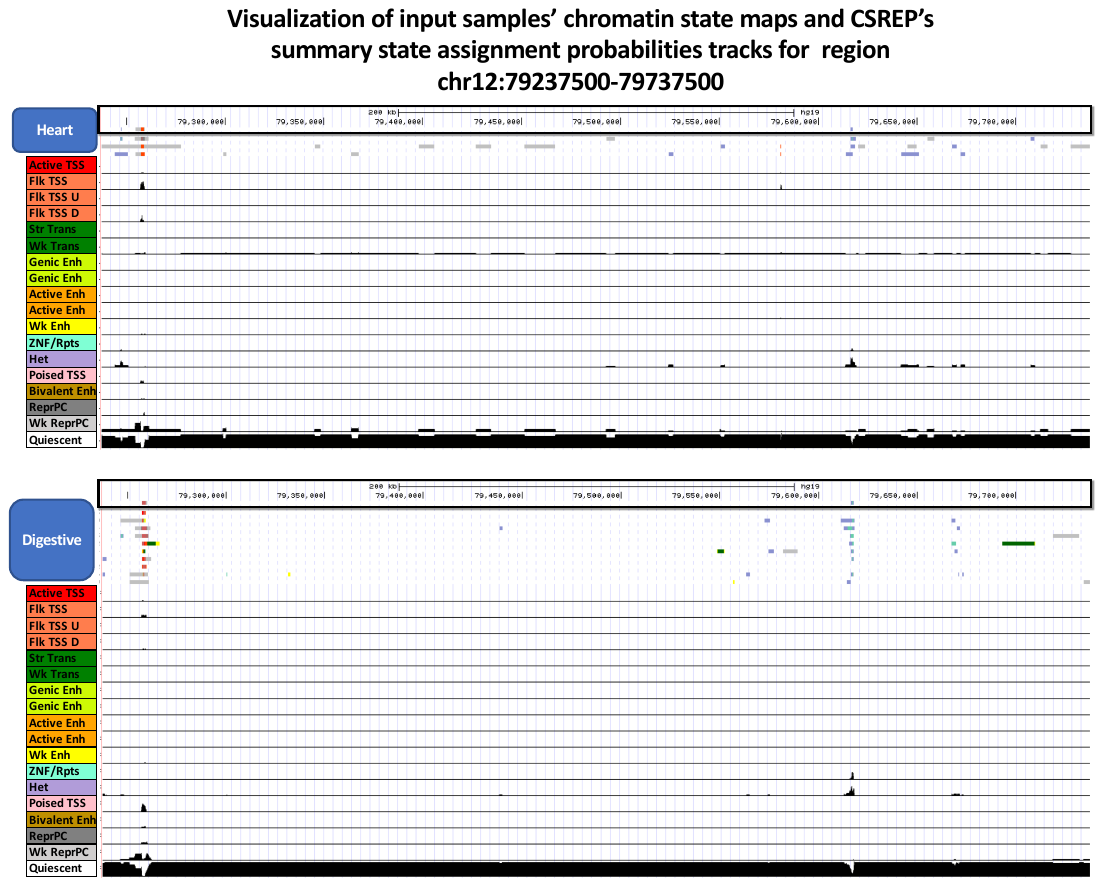

**Supplementary Figure 2**: **Visualization of CSREP’s input and output data for an arbitrary 500-kb genomic window (chr12:79,237,500-79,737,500, hg19).** Similar to Supp. Fig. 1, for genomic region chr12:79,237,500-79,737,500, hg19.

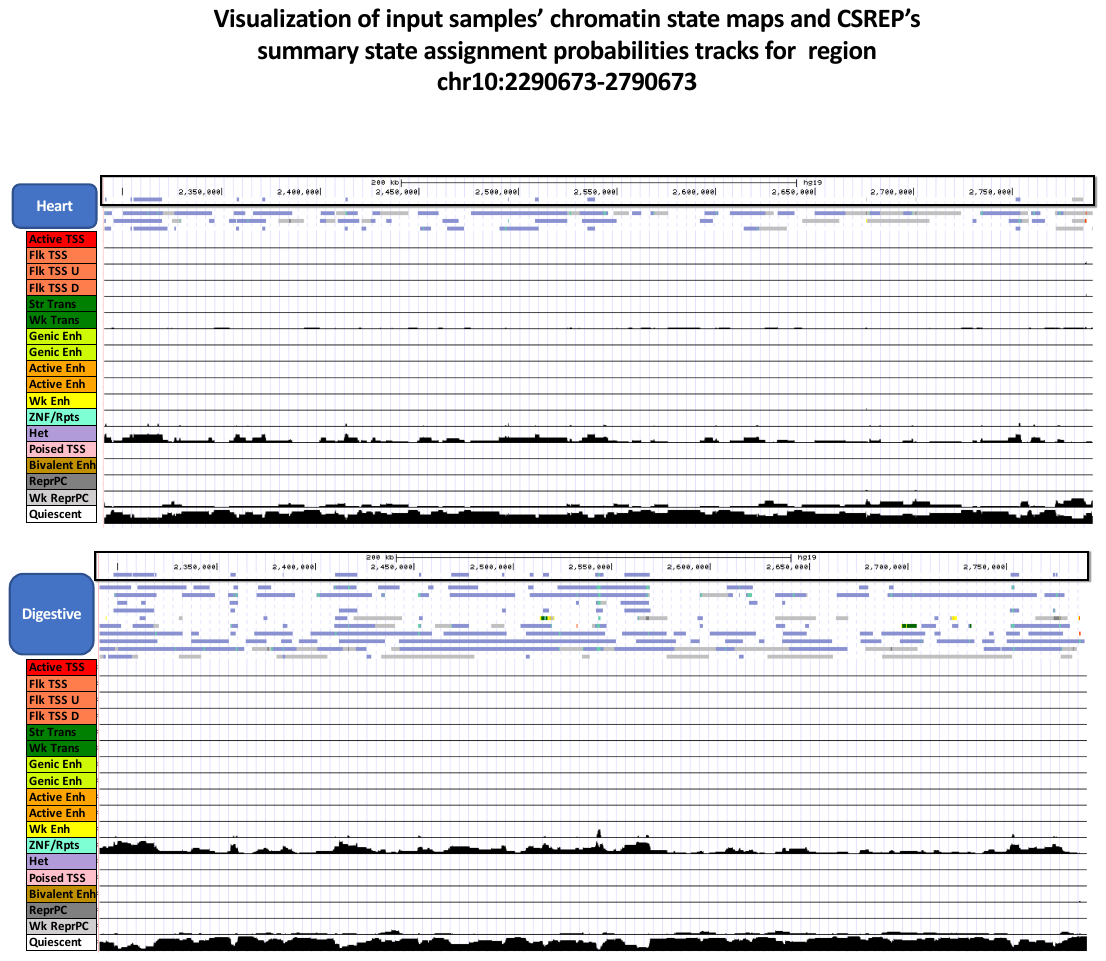

**Supplementary Figure 3**: **Visualization of CSREP’s input and output data for an arbitrary 500-kb genomic window (chr10:2,290,673-2,790,673, hg19).** Similar to Supp. Fig. 1, for genomic region chr10:2,290,673-2,790,673, hg19.

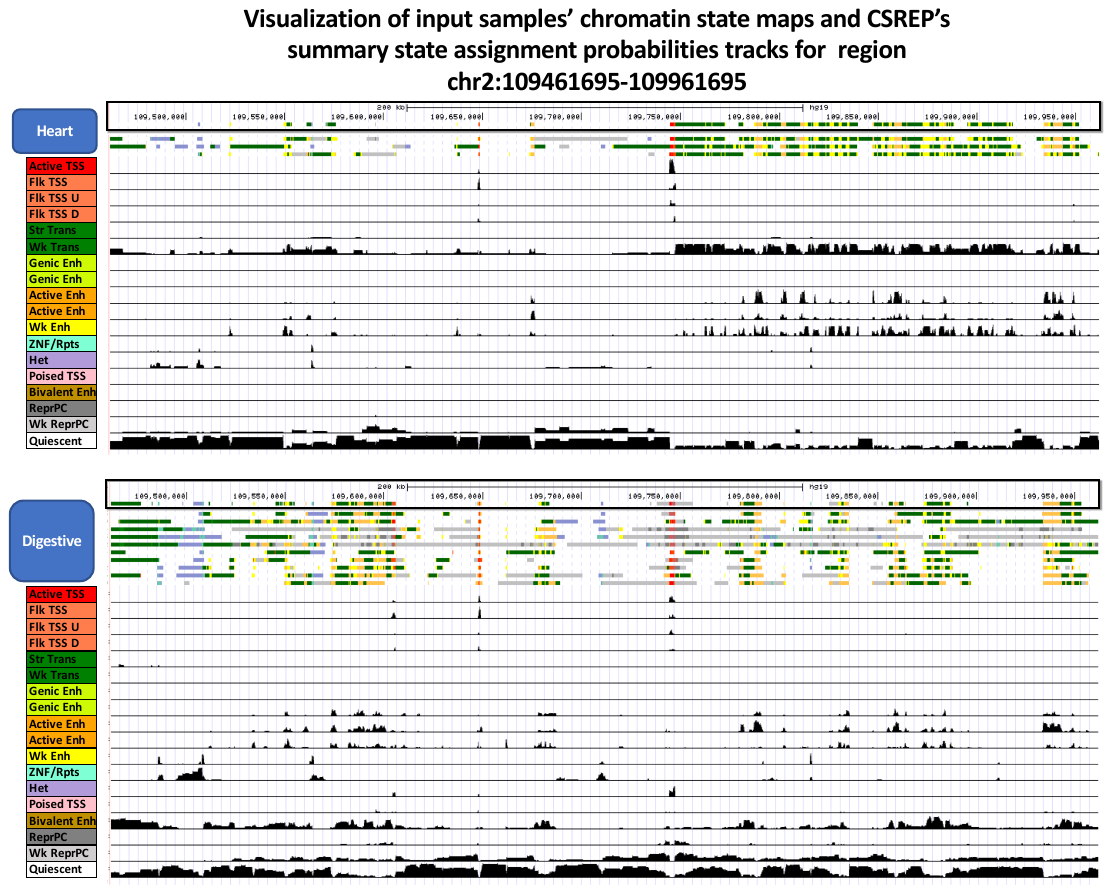

**Supplementary Figure 4**: **Visualization of CSREP’s input and output data for an arbitrary 500-kb genomic window (**chr2:109,461,695-109,961,695, hg19**).** Similar to Supp. Fig. 1, for genomic region chr2:109,461,695-109,961,695, hg19.

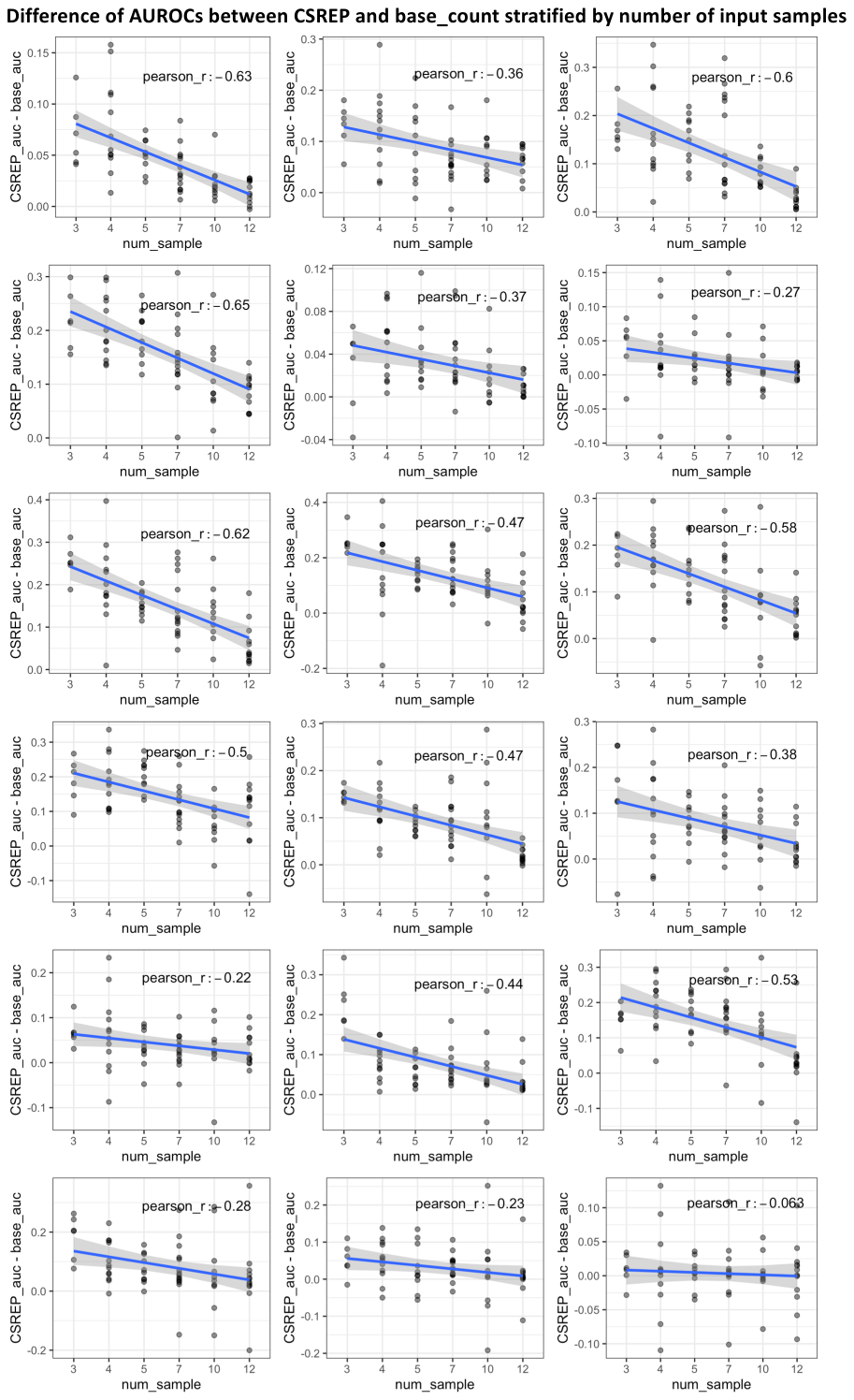

**Supplementary Figure 5:** **Relationship between the number of samples and AUROCs from using summary chromatin state map to predict genomic locations of individual chromatin states**. We conducted cross-validation analysis for each group of samples (**Supp. Methods**), and for each group, we calculate ROC curve of CSREP’s summary probabilistic chromatin state map in recovering genomic positions of individual chromatin states in a held-out sample. Each panel corresponds to a chromatin state, and shows the difference between CSREP’s AUROCs and base_count’s AUROCs for predicting locations of the chromatin state in left-out samples. Each dot corresponds to one sample. Y-axis shows the difference of AUROCs between the two methods (positive y-axis means CSREP results in higher AUROCs and vice-versa). X-axis shows the number of input samples for the group.

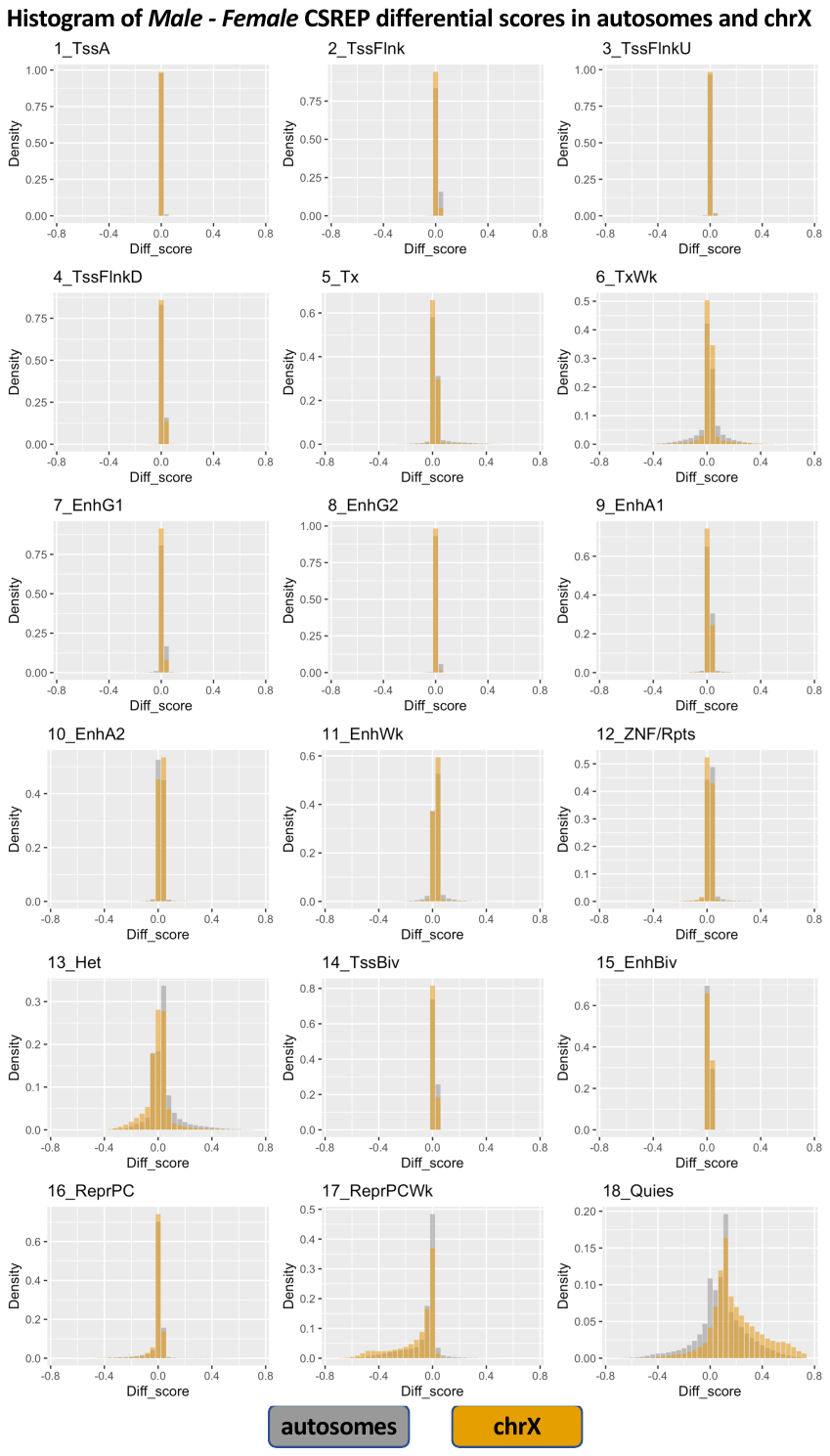

**Supplementary Figure** **6**: **Histogram of CSREP differential chromatin scores between Male and Female groups of samples, in autosomes and in chromosome X.** Each subpanel shows the histograms of one state’s CSREP *Male - Female* differential scores, bounded between -1 and 1, in autosomes and chromosome X.

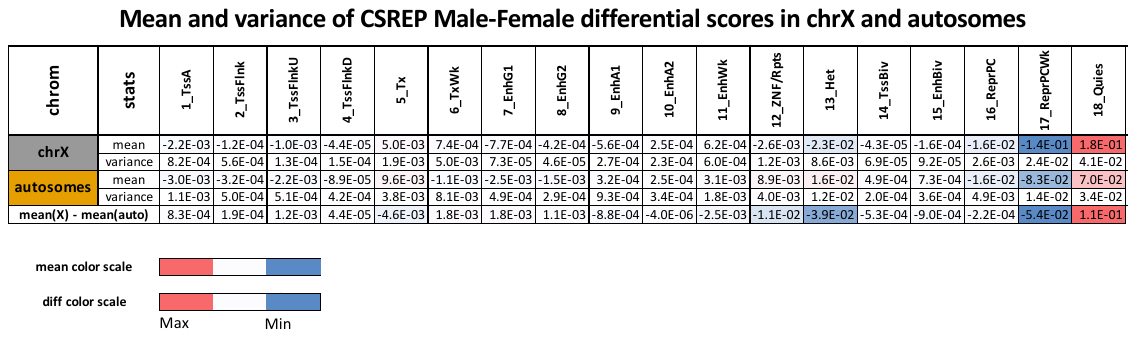

**Supplementary Figure 7:** **Mean and variance of the CSREP differential scores between Male and Female groups of samples, in autosomes and in chromosomes.** The mean differential scores for each state in either chromosome X or autosomes are reported for each state and share the same color scale as in bottom legend. The difference between the mean scores for each state is reported on the bottom row and colored as in bottom legend. Three states with largest-magnitude difference in mean scores are 13_Het, 17_ReprPCWk, 18_Quies.

**References**

Boix,C.A. *et al.* (2021) Regulatory genomic circuitry of human disease loci by integrative epigenomics. *Nature*, **590**, 300–307.

Ebert,P. and Schulz,M.H. (2020) Fast detection of differential chromatin domains with SCIDDO. *Bioinformatics*.

Roadmap Epigenomics Consortium,A. *et al.* (2015) Integrative analysis of 111 reference human epigenomes. *Nature*, **518**, 317–330.

Quinlan,A.R. and Hall,I.M. (2010) BEDTools: a flexible suite of utilities for comparing genomic features. *Bioinformatics*, **26**, 841–842.
